# Bacterial keratitis: similar bacterial and clinical outcomes in female versus male New Zealand White rabbits infected with *Serratia marcescens*

**DOI:** 10.1101/2021.08.25.457638

**Authors:** Eric G. Romanowski, Sanya Yadav, Nicholas A. Stella, Kathleen A. Yates, John E. Romanowski, Deepinder K. Dhaliwal, Robert M. Q. Shanks

## Abstract

Females and males respond differently to a number of systemic viral infections. Differences between females and males with respect to the severity of keratitis caused by Gram-negative bacteria such as *Serratia marcescens* are less well established. In this study we injected female and male New Zealand White rabbit corneas with a keratitis isolate of *S. marcescens* and evaluated the eyes after 48 hours for a number of clinical and microbiological parameters. No statistical differences in bacterial burden and corneal scores were recorded between female and male rabbits although there was a non-significant trend toward a higher frequency of female rabbits demonstrating hypopyons. This data suggests that for experimental bacterial keratitis studies involving Gram-negative rods, a single sex or mixed group of rabbit is sufficient for evaluating pathology and bacterial burdens. This will reduce the number of animals used for subsequent studies.

## 1. INTRODUCTION

Microbial keratitis is a vision threatening inflammatory condition of the cornea resulting from an underlying infection caused by bacteria, viruses, fungi, and acanthamoeba. It has been estimated that 930,000 doctor’s office and outpatient clinic visits and 58,000 emergency department visits occur annually for keratitis or contact lens problems [1]. Some prior studies provide cumulative data on the incidence of keratitis and outcomes of microbial keratitis, but they lack analyses of sex differences among the patients [1–4].

Few reports demonstrate sex-based difference in patients with eye infections. Krishnan et al. showed that the corneas of female fungal corneal ulcer patients took longer to re-epithelialize compared to males, but ultimately there were no differences in visual acuity or infiltrate/scar size [5]. Sagara and colleagues reported women were more prone to bleb-related infections after glaucoma filtering surgery in the spring season in Japan [6]. Previous studies have found men to be at a higher risk of acanthamoeba keratitis due to a number of factors including being less compliant with contact lens care and for a variety of other hygiene behaviors [7]. Sex differences are known to exist for immune response to systemic viral infections [8] as well as to their prophylaxis and therapy [9] per infectious disease literature. However, data regarding sex differences in ocular viral keratitis, which are mainly based on animal models, are mixed. One study reported no significant sex differences in ocular HSV-1 infection [10], whereas another study found consistent worse outcome in males [11]. With regards to the ocular microbiome, sex differences were demonstrated in the bacterial species making up the conjunctival microbiome among non-contact lens wearers [12].

Few studies have directly explored the role of sex in bacterial keratitis. Some epidemiological studies on bacterial keratitis have reported no significant sex related differences in clinical presentation and outcomes among patients [13, 14]. In one study investigating clinical records of patients with *Serratia marcescens* at Wills Eye Hospital, sex was not directly indicated to play a role [15]. However, in a nationwide study of all infections caused by *S. marcescens* in Canada, males were infected at a significantly higher rate [16]. Although, sex differences for ocular infections were not tested, this study suggested that sex could play a role in clinical outcomes of bacterial keratitis caused by *S. marcescens*.

The dearth of studies evaluating sex difference in patients with bacterial keratitis led us to the current study, in which we investigated sex as a biological variable in clinical presentations and outcomes in male and female New Zealand white (NZW) rabbit corneas infected with the Gram-negative bacterium *S. marcescens*. This study could inform future experimental bacterial keratitis studies involving Gram-negative rods, that a single sex of rabbit is sufficient for evaluating pathology and bacterial burdens. This will ultimately reduce the number of animals used for subsequent studies.

## 2. MATERIALS AND METHODS

### 2.1. Bacterial strain

*S. marcescens* strain K904 is a strain isolated from a case of contact lens related bacterial keratitis at the Charles T. Campbell Ophthalmic Microbiology Laboratory at the UPMC Eye Center in the Department of Ophthalmology at the University of Pittsburgh School of Medicine. Its characteristics have been previously described [17].

### 2.2. Animals

Female and male New Zealand white (NZW) rabbits weighing 1.1-1.4 kg were obtained from the Oakwood Research Facility (Oxford, MI) through Charles River Laboratories while some male animals were sourced from Charles River’s Canadian rabbitry. In addition, 5 female and 3 male NZW rabbits weighing 1.2-1.35 kg were donated by the Department of Laboratory Animal Resources at the University of Pittsburgh. These rabbits were the result of an unintended breeding. The dam was originally purchased from Covance Research Products (Denver, PA). The current study conformed to the ARVO Statement on the Use of Animals in Ophthalmic and Vision Research and was approved by the University of Pittsburgh’s Institutional Animal Care and Use Committee (IACUC Protocols 16098925 and 19085820).

### 2.3. Bacterial keratitis studies

For the keratitis studies, *S. marcescens* strain K904 was grown overnight at 30°C with aeration, normalized by optical density (600 nm), and adjusted to ~1000 CFU in 25 μl in PBS. The rabbits were systemically anesthetized with 40 mg/kg of ketamine and 4 mg/kg of xylazine administered intramuscularly. The corneas of the right eyes only were anesthetized with topical 0.5% proparacaine and injected intrastromally with the 25 μl of *S. marcescens*. The actual inoculum colony counts for each trial were determined using the EddyJet 2 spiral plating system (Neutec Group Inc., Farmingdale, NY) on 5% trypticase soy agar with 5% sheep’s blood plates. The plates were incubated overnight at 30°C and the colonies were counted using the automated Flash and Grow colony counting system (Neutec Group, Inc).

At 48 h post-injection, the eyes were evaluated for ocular signs of inflammation using a slit-lamp according to a modified McDonald-Shadduck grading system [18]. Following the 48 hour examination, the rabbits were euthanized with an intravenous overdose of Euthasol Solution following systemic anesthesia with ketamine and xylazine. Subsequently, corneal buttons were harvested using a 10 mm trephine and placed into Lysing Matrix A tubes (MP Biomedicals) containing 1 ml of PBS. The corneas were then homogenized with an MP Fast Prep-24 homogenizer (MP Biomedicals), and the numbers of corneal bacteria were enumerated as described above.

### 2.4. Statistical analysis

Data was analyzed using GraphPad Prism Software. Non-parametric analysis was used to analyze inflammatory scores by Mann-Whitney or Kruskal-Wallis analysis, and ANOVA with Tukey’s post-test or Student’s T-tests were used to analyze corneal colony count data.

## 3. RESULTS

### 3.1. Bacterial burden during *S. marcescens* keratitis was similar in female and male rabbits

Female and male rabbits were injected in the corneal stroma with approximately 1000 CFU of a wild-type contact lens isolate of *S. marcescens*, strain K904 (Figure 1A). At 48 h post-injection corneas were harvested and the bacterial burdens within the corneas were enumerated by plating dilutions of the corneal homogenate on blood agar plates. Over 10^7^ CFU were found in both male and female corneas with no statistical difference (p=0.772) (Figure 1B). However, the median number of bacteria isolated from corneas was slightly higher in females (7.50 Log_10_ CFU, CI 6.80-7.76) than in males (7.30 Log_10_ CFU, CI 6.31-7.66).

**Figure 1.**
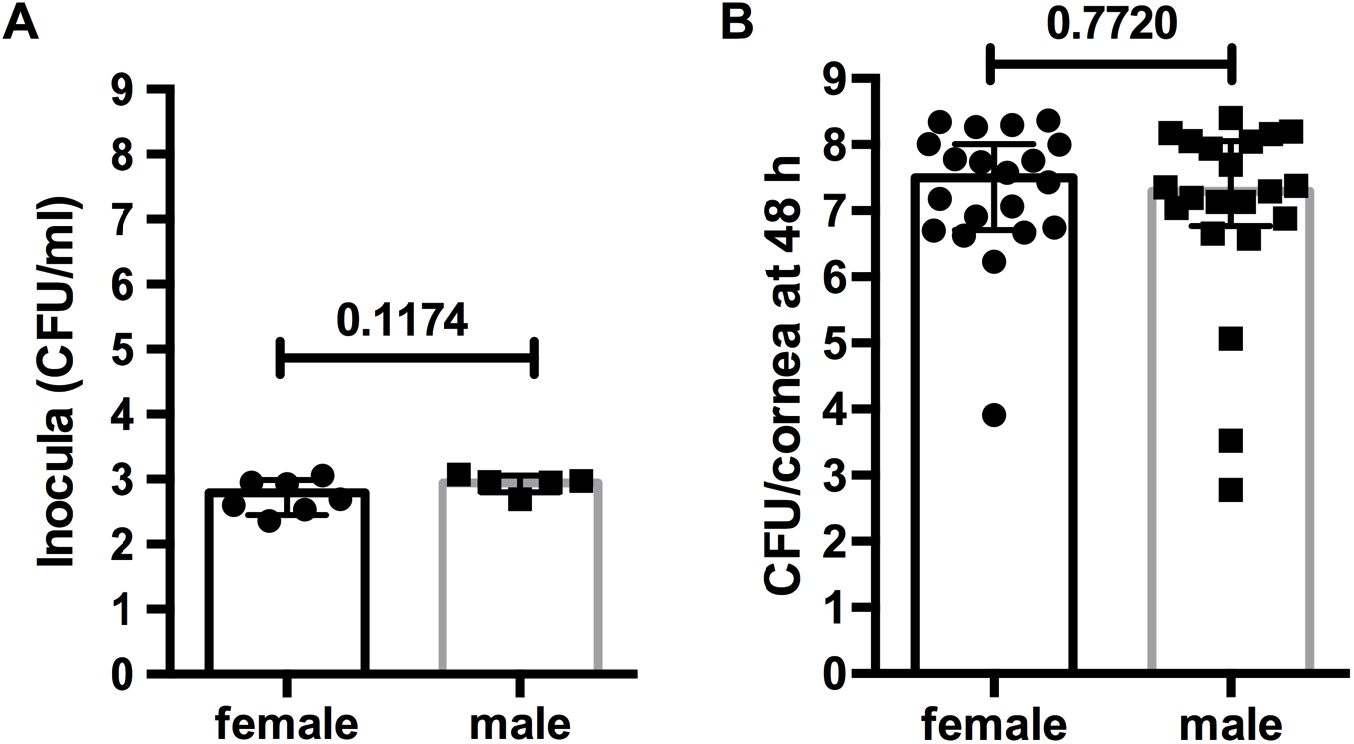
Bacterial colony counts of inocula and infectious burdens. The pre-injection inocula **(A)** and the infectious burdens from homogenized corneas from 48 h post infection **(B)** were determined following growth on blood agar plates. The medians and interquartile ranges are shown, p-values were determined using the Mann-Whitney test.

### 3.2. The clinical presentation of corneal inflammation was similar between female and male rabbits infected by *S. marcescens* strain K904

Rabbit corneas infected by *S. marcescens* generally exhibited corneal infiltrates with a snowflake-like morphology and mild iritis at 24 h in both sexes, similar to a previous report [19] (Figure 2). At 48 hours, the pathology of the eyes had clearly progressed. The eyes presented with severe purulent discharge, and the corneas demonstrated diffuse haziness around large, dense, central infiltrates with surrounding rings, a hallmark of Gram-negative bacteria related keratitis [20] in both sexes. Intraocularly, the eyes demonstrated inflammation manifested by moderate iritis, diffuse fibrin production and accumulations of fibrin or hypopyons in the anterior chamber of most animals from both sexes. Staining with fluorescein revealed ulcers above the infiltrates in most cases (Figure 2).

**Figure 2.**
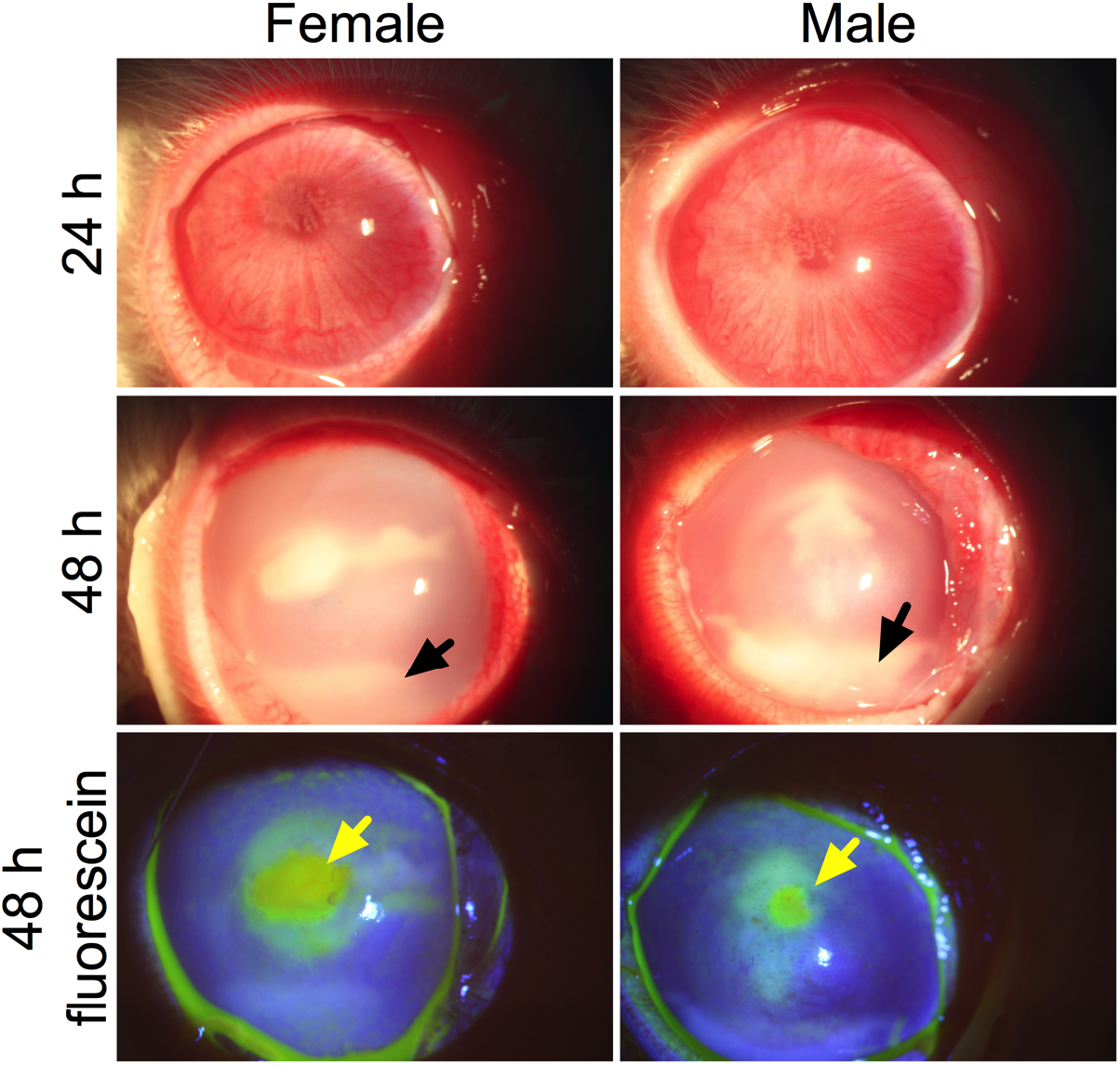
Rabbit eyes at 24 and 48 h post infection with *S. marcescens*. Representative images of eyes with median corneal inflammatory scores are depicted. Black arrows indicate the presence of a hypopyon in the anterior chamber and yellow arrows indicate corneal epithelial erosions stained by fluorescein.

Corneal scores, based on the MacDonald-Shadduck test [18], were identical between females and males with a median score of 2.0 at 24 h and 8.0 at 48 h for both, p>0.0.5 (Figure 3A-B), whereas PBS injected control eyes had a median score of zero (n=14). Corneal defect areas were not significantly different between sexes at 48 h (p=0.7394), although the median was slightly higher for females 5.0 mm^2^ versus males 3.1 mm^2^ (Figure 3C). However, there was more variation among male eyes with respect to erosions and the mean value was higher for males (7.8 ± 11.4 mm^2^ versus 5.2 ± 4.8 mm^2^ for females). The presence of eyes with fibrin or hypopyons in the anterior chamber was greater for females (100%) than males (85.7%); however, the difference was not statistically significant (p=0.0820).

**Figure 3.**
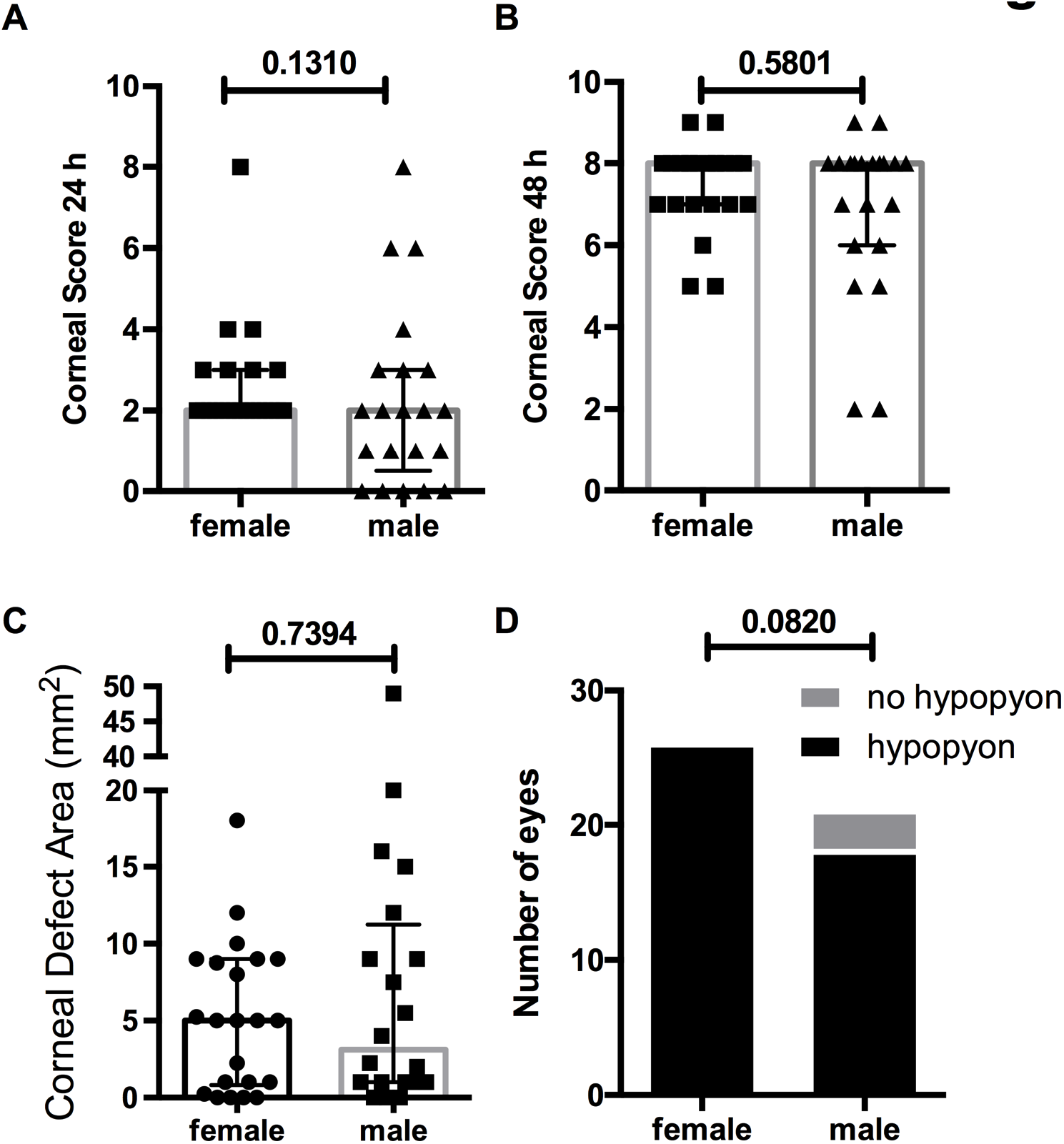
Pathology of infected corneas. Clinical inflammation was assessed at 24 **(A)** and 48 h post infection **(B)** using a modified McDonald-Shadduck scoring system. The area of corneal epithelial erosion was determined **(C)**. The number of eyes demonstrating a hypopyon or fibrin in the anterior chamber were counted **(D)**. The medians and interquartile ranges are shown **(A-C)**. p-values were determined using the Mann-Whitney test **(A-C)** or Fisher’s exact test **(D).**

## 4. DISCUSSION

The National Institutes of Health research grant proposals require the use of both female and male animals unless appropriate justification is provided. In many cases it is essential to use animals of both sexes due to differences in disease outcomes [21, 22]. In other situations, where there are no major differences between female and male animal models [10, 23], The use of both sexes leads to the unnecessary use of animals. This is especially an issue with models that require Animal Welfare Act covered animals, such as rabbits.

With the exception of *Pseudomonas aeruginosa*, it is difficult for most bacteria that infect human corneas to create similar infections with rodent models. To get around this, existing models use mutant mice with incomplete immune systems. However, these do not fully replicate the natural host-pathogen interactions either. Rabbit eyes, on the other hand are well suited for the purpose, as they are readily infected by major ocular pathogens such as staphylococci and Enterobacteriales. However, rabbits are expensive. Moreover, one of the major tenants of the 3Rs [24] is to reduce animal numbers in experimental studies. For addressing these issues, we performed this study to evaluate whether we needed to include both sexes when analyzing bacterial keratitis.

We observed that that were no significant pathological differences between female and male New Zealand rabbit corneas infected by a representative strain of *S. marcescens*, the Enterobacterales species most frequently responsible for causing bacterial keratitis [25]. Numerous parameters in pathogenesis and clinical outcomes were evaluated, and while there was a slight trend toward more severe disease in females, the difference was minor and not significant. Therefore, we conclude that future work using *S. marcescens* does not justify the use of both sexes, and that one sex or a mix is sufficient. This is likely the same for other Gram-negative rods, but this remains to be formally demonstrated by future studies. Studies such as this one can help researchers to eliminate unnecessary expenses by reducing animal numbers.

## Acknowledgements

This work was funded by National Institute of Health grants EY027331 to R.M.Q.S and Core Grant for Vision Research EY08098. Additional funds for the department were provided by the Eye and Ear Foundation of Pittsburgh and unrestricted funds from Research to Prevent Blindness.

